# Analysis of the effects of mating systems on lineage diversification across multiple genera

**DOI:** 10.1101/2024.08.20.608795

**Authors:** Kuangyi Xu

**Affiliations:** Department of Ecology and Evolutionary Biology, University of Toronto, 25 Willcocks St, Toronto, ON, Canada M5S 3B2

**Keywords:** mating system, speciation, diversification, selfing, life form, hidden state, mixed mating

## Abstract

- The transition from outcrossing to self-fertilization is a major evolutionary trend in plants, with selfing long hypothesized an evolutionary dead end. Recent theories suggest that elevated extinction rates may be restricted to highly selfing rather than mixed-mating populations. Previous analyses of the effects of mating systems on diversification found mixed results, varying by focal clades and potentially suffering from several limitations.
- We collected data on three mating-system-related characters. We also collected life form data to indirectly distinguish between mixed mating and highly selfing. We estimated speciation and extinction rates for character states, and state transition rates in 27 genera.
- We find that outcrossing lineages diversify at higher rates than selfing lineages, while the impact of mating systems on speciation is not consistent across genera. Among self-compatible lineages, annuals/short-lived exhibit significantly lower diversification rates than perennials. Models incorporating hidden states often provide a better fit than those without.
- The results suggest that selfing lineages overall have lower diversification rates. This may be primarily contributed by elevated extinction rates in highly selfing lineages, which may also explain the prevalence of mixed-mating systems. However, the influences of mating systems on diversification may be often driven by other factors correlated with mating system transitions.

## Introduction

The transition from outcrossing to self-fertilization is one of the major evolutionary trends in flowering plants (Barrett, 2008), with selfing occurring in about 50% of species (Goodwillie et al., 2005). However, it has long been hypothesized that selfing is an evolutionary dead end (Igic & Busch, 2013). Transitions from selfing back to outcrossing are rarely observed in nature (Barrett, 2013; Fujii et al., 2016), and self-incompatibility is hard to be regained once it is lost (Igic et al., 2008). Compared to outcrossing lineages, selfing lineages are thought to suffer higher extinction risks due to several microevolutionary effects of selfing (Hartfield et al., 2017).

Selfing increases homozygosity in the offspring, and reduces the effective population size due to larger variance in the number of alleles an individual leaves to the next generation (Caballero & Hill, 1992). Increased genome-wide homozygosity also reduces effective recombination rates, which, in the presence of deleterious mutations, can lead to a further reduction of the effective population size (background selection; Nordborg et al., 1996) and the accumulation of deleterious mutations (Charlesworth et al., 1993; Xu, 2022). Moreover, theoretical work and meta-analyses suggest that selfers tend to exhibit lower levels of additive genetic variance in quantitative traits compared to outcrossers (Clo et al., 2019), although selfing may temporarily reveal non-additive genetic variance in the short term (Clo et al., 2020; Clo & Opedal, 2021).

Due to the reduced efficacy of selection in selfing populations, selfers are expected to exhibit a higher ratio of nonsynonymous to synonymous divergence (*D*_*n*_/*D*_*s*_) or polymorphism (*π*_*n*_/*π*_*s*_) compared to outcrossers (Glémin, 2007; Wright et al., 2008). Some studies did not detect significant differences in these ratios when comparing sister species with contrasting mating systems (Escobar et al., 2010). This is often attributed to the relatively recent origin of selfing lineages, where inbreeding may not have persisted long enough to generate significant differences. More recent studies, however, have found higher nonsynonymous-to-synonymous ratios in selfers (Ness et al., 2012; Arunkumar et al., 2015; Wang et al., 2021).

The impacts of selfing on speciation remain poorly understood, as speciation can occur through various processes (Coyne & Orr, 2004), and selfing can influence them in complex ways (Marie-Orleach et al., 2024). Selfing may reduce gene flow between populations and contribute to reinforcement (Castillo et al., 2016), but can also hinder the accumulation of alleles that drive speciation by reducing effective population size and effective recombination rates. Recent studies suggest that Dobzhansky-Muller incompatibilities might accumulate more rapidly in selfing populations (Marie-Orleach et al., 2022; Marie-Orleach et al., 2024). Additionally, transitions to selfing are often associated with a series of morphological changes (selfing syndrome; Sicard & Lenhard, 2011), potentially contributing to reproductive isolation (Cutter, 2019). Furthermore, selfing may facilitate polyploid speciation by allowing polyploid species to avoid mating with diploid individuals during the early stages of establishment (Rausch & Morgan, 2005; Barringer, 2007).

The macroevolutionary effects of selfing on lineage diversification have been estimated using state-dependent speciation and extinction (SSE) models (Helmstetter et al., 2023), but the results vary across clades. Several studies have found that lineages with traits promoting selfing tend to exhibit reduced rates of net diversification (speciation minus extinction) (Goldberg et al., 2010; de Vos et al., 2014; Gamisch et al., 2015; Castillo et al., 2016; Freyman & Höhna, 2019). However, opposite results have been observed in other clades (Castillo et al., 2016; Landis et al., 2018). These studies also found mixed results for the effects of selfing on speciation, with selfing enhancing speciation rates in some clades while inhibiting speciation in others.

Several potential limitations exist in previous diversification analyses. Most of these studies were conducted at the family level, which may bias results toward genera with more complete character state records. For instance, in the analyses of Solanaceae and Onagraceae, over two-thirds and one-third of the included species are from *Solanum* and *Oenothera*, respectively (Goldberg et al., 2010; Freyman & Höhna, 2019). Additionally, in many family-level analyses, mating system polymorphism is primarily attributed to variation between genera, while congeneric species often share identical mating system states. This can result in estimates being confounded by unknown genus-specific traits. In fact, SSE models are prone to inferring false positives; even if diversification rate variation is driven by other factors, a significant dependency on the focal character may still be inferred if state transitions coincide with more or less diverse parts of the tree (Rabosky & Goldberg, 2015). For example, the breakdown of self-incompatibility may be often triggered by unfavorable environmental shifts, such as pollen limitation (Busch & Schoen, 2008; Eckert et al., 2010; Encinas-Viso et al. 2020). In this case, it may be pollen limitation, rather than the breakdown of self-incompatibility, that drives extinction.

The problem of coincidental associations and clade dependency may be mitigated by comparative analyses across multiple clades. For example, Sabath et al. (2016) showed that dioecy neither consistently inhibits nor facilitates diversification across more than 30 genera. Additionally, the issue of false positives has been explicitly treated by recently developed methods, such as the HiSSE model (Hidden State Speciation and Extinction; Beaulieu & O’Meara, 2016). HiSSE models extend BiSSE models by allowing speciation and extinction rates to depend on both measured and hidden character states. HiSSE analysis in *Onagraceae* found that, for all hidden states, self-compatible lineages consistently exhibit lower diversification rates than self-incompatible lineages (Freyman & Höhna, 2019). However, HiSSE analysis in Polemoniaceae found that diversification rates appear to be independent of whether lineages are autogamous or cross-pollinated (Landis et al., 2018).

Importantly, early theoretical and empirical investigations often focus on differences between predominantly outcrossing and predominantly selfing populations, while recent progresses suggest that the rate of selfing is critical. Specially, a higher selfing rate reduces the fixation probability of new adaptive mutations at a single locus only when alleles are dominant; however, very high selfing rates significantly strengthens background selection, thereby inhibiting fixation even for recessive alleles (Glémin & Ronfort, 2013). Additionally, intermediate selfing rates are optimal in promoting rescue through the sweep of adaptive alleles at multiple loci by preventing recombination from destroying haplotypes with multiple adaptive alleles (Xu, 2023a). Populations with intermediate selfing rates may also avoid the rapid accumulation of deleterious mutations that occurs in highly selfing populations (Xu, 2022). This is because even a slight amount of effective recombination introduced by outcrossing is sufficient to significantly slow down Muller’s ratchet. Furthermore, compared to outcrossing populations, populations with intermediate selfing rates tend to have lower genetic load and are more likely to persist under pollen limitation (Busch & Delph, 2012; Roze, 2015; Xu, 2023b).

The above results suggest the hypothesis that elevated extinction rates may be restricted to highly selfing, instead of intermediate selfing populations. This could partly explain why intermediate selfing is common in nature, despite being predicted to be evolutionarily unstable, while highly selfing, which is predicted to be evolutionarily stable, seldom occurs (Goodwillie et al., 2005). Specifically, under the balance between lineage diversification and mating system transitions, the much lower diversification rates in highly selfing lineages compared to mixed-mating lineages could result in a low prevalence of highly selfing species and a substantial proportion of mixed-mating lineages found in nature.

However, previous diversification analyses treated mating systems as binary-state characters (e.g., self-incompatible/self-compatible). The binary-state classification provides no information about the rate of selfing, since being self-compatible does not necessarily mean selfing occurs at a high rate (Raduski et al., 2012). Therefore, testing the proposed hypothesis necessitates diversification analyses that differentiate among predominantly outcrossing, mixed mating, and highly selfing lineages.

To mitigate the limitations of previous analyses and clade dependency, we revisit the question of the macroevolutionary consequences of selfing. We conducted BiSSE analyses across 27 genera using a Bayesian framework. To reduce false positives, we also applied HiSSE models to these genera and examined the effects of mating systems on diversification rates across different hidden states.

To test the hypothesis that highly selfing lineages tend to have lower diversification rates than those with intermediate selfing rates, it would be ideal to conduct diversification analyses based on outcrossing rates. However, we did not identify appropriate genera with sufficient data for this analysis. Therefore, we performed diversification analyses on the combination of self-compatibility and life form states (annual/short-lived *versus* perennial). Given that annuals are much more likely to be highly selfing compared to perennials (Aarssen, 2000; Ma et al., 2020), we expect that self-compatible annuals will generally exhibit lower diversification rates and higher extinction rates than self-compatible perennials.

## Materials and methods

### Data collection

We collected datasets for three characters related to the mating system: self-compatibility, pollination system, and species-level outcrossing rate. The genus name were updated based on the WFO Plant List (https://wfoplantlist.org). For the pollination system, we directly used the dataset reported from Landis et al. (2018). The self-compatibility dataset was primarily obtained by compiling datasets reported in previous works (Wagner et al., 2017; Grossenbacher et al., 2017; Freyman & Höhna, 2019; Delaney & Igić, 2022; Krakos et al., 2022; Cascante-Marín et al., 2023). For the dataset reported by Delaney & Igić (2022), we only included self-compatibility data based on hand-pollination experiments, while sources marked as verbal descriptions without experimental validation were considered unreliable and not used. In addition, we included self-compatibility data for a few species that were previously missing through a literature search. A species is classified as self-incompatible (SI) when the index of self-compatibility is less than 0.2; otherwise, it is considered self-compatible (SC). In cases where self-compatibility was measured or reported for multiple populations of a species, we determined self-compatibility at the species level by selecting the self-compatibility status that occurred in the majority of conspecific populations.

For the outcrossing rate dataset, we assembled data from three previous studies (Dumini et al., 2009; Goodwillie et al., 2012; Whitehead et al., 2018). We also added estimations from more recent publications. To do this, we focused exclusively on multi-locus estimates of outcrossing rates, as these estimates are less influenced by selection compared to single-locus estimates (Ritland, 2002; Bürkli et al., 2017). To identify relevant studies, we searched the Web of Science up to November 12, 2023, for publications that cited at least one of five methodological papers on multi-locus outcrossing rate estimations (Ritland, 1990; Ritland, 2002; Ritland & Jain, 1981; David et al., 2007; Jones & Wang, 2010). For a species, when outcrossing rate estimates were available for multiple populations or multiple flowering seasons, we averaged these estimates to calculate a mean outcrossing rate at the species level. We recognized that outcrossing rates often vary among conspecific populations. However, meta-analyses indicate that this variation tends to be small in predominantly outcrossing and predominantly selfing species (Whitehead et al., 2018), suggesting it may be valid to classify the mating system of a species into discrete categories.

We classified the mating-system-related characters into binary states, with 0 representing outcrossing and 1 representing selfing. Specifically, for self-compatibility, state 0 indicates self-incompatible (SI) species, and state 1 indicates self-compatible (SC) species. For the pollination system, state 0 and 1 correspond to cross-pollination and autogamy, respectively. For outcrossing rates, state 0 represents species with mean outcrossing rates above 0.8, and state 1 represents species with mean outcrossing rates below 0.8.

Two genera, *Solanum* and *Oenothera*, were found to be suitable to conduct diversification analyses on the combination of self-compatibility and life form states. The life forms of *Solanum* species were primarily determined using the dataset from Hilgenhof et al. (2023). Species marked as shrub or tree in Hilgenhof et al. (2023) were classified as perennials. For species marked as herb, we determined whether they are short-lived or perennial primarily based on descriptions from Solanaceae Source (http://solanaceaesource.org) with additional literature searching. Life forms of *Oenothera* species were determined based on descriptions in Dietrich & Wagner (1988), Wagner et al. (2007), and Evans et al. (2011).

### Phylogeny reconstruction

For genera selected from the pollination system dataset, we directly used the maximum likelihood (ML) trees and bootstrap trees reported by Landis et al. (2018). For other genera, phylogenies were reconstructed following the procedure described below.

1. For each genus, we selected outgroup species based on prior phylogenetic studies (summarized in Table S1). Sequence data for all species within the genus, as well as the outgroup species, were downloaded from GenBank and clustered using the R package *phylotaR* (Bennett et al., 2018).
2. Within each cluster, sequences that were either twice as long or half as short as the median sequence length were discarded. The best-represented sequence for each species was selected using the “drop_by_rank” function in the *phylotaR* package with default settings. Sequences within each cluster were then aligned using MAFFT version 7 (Katoh & Standley, 2013), and unreliable sites were removed using Gblocks (Castresana, 2000). Clusters containing sequences from only few species were dropped. However, since previous studies suggest that filtering genes based on missing data may reduce phylogenetic accuracy (Molloy & Warnow, 2018), most clusters were retained. The remaining clusters were concatenated into a partitioned supermatrix for phylogenetic reconstruction.
3. For each genus, a ML tree and 100 bootstrap trees were reconstructed using raxmlGUI 2.0 (Edler et al., 2021) with an initial model test to select the best-fit model. To ultrametricize the ML and 100 bootstrap trees, we employed the “chronos” function from the R package *ape* (Paradis & Schliep, 2019). The smoothing parameter *λ* in the “chronos” function, which adjusts the penalty for rate variation across branches (Sanderson, 2002), was determined through cross-validation on the ML tree. The cross-validation was conducted using the “CV” function from the R package *chronos* (https://github.com/josephwb/chronos), varying *λ* from 0.01 to 1000 at intervals of 0.5 on a log10 scale. After ultrametricization, outgroups and species with missing character state information were pruned for diversification analyses.

To reduce uncertainty caused by small sample sizes, we excluded genera from the diversification analyses if their resulting phylogenies contained fewer than nine species or if there were fewer than three species in each character state, unless the total number of species in the genus was small. As a low sampling proportion reduces the confidence of SSE analyses (Helmstetter et al., 2023), genera with excessively sampling fractions (i.e., the proportion of species with character state data relative to the total number of species) lower than 5% were also excluded from the analyses. Mating system character states, supermatrices, and phylogenies for genera used in our diversification analysis are available at https://doi.org/10.5281/zenodo.13265323. The number of species in each mating system state and summary statistics of supermatrices for genera used in our diversification analyses are available at Table S2.

### Diversification analyses

We assessed the impacts of mating system states on lineage diversification using several SSE models. To account for phylogenetic uncertainty, diversification analyses were performed on 100 bootstrap trees for each genus. Briefly, we estimated diversification for outcrossing (state 0) and selfing (state 1) lineages using the BiSSE models (Maddison et al., 2007), implemented in the R package *diversitree* (Fitzjohn, 2012). We also fitted a series of HiSSE models that include hidden states using the R package *hisse* (Beaulieu & O’Meara, 2016). For genera with sufficient character data, we additionally estimated diversification rates for the combination of self-compatibility and life form states using MuSSE models (Multiple State Speciation and Extinction), implemented in the *diversitree* package. Since estimations of extinction rates were suggested to be unreliable (Rabosky, 2010), our analyses focused on the speciation and diversification rates (speciation minus extinction).

All the SSE analyses were conducted by accounting for the sampling fraction of species included in the analyses relative to the total number of species in a genus, which was determined using the number of accepted species in the WFO Plant List. Since we lacked complete character state information for all taxa in each genus, we assumed no state-dependent sampling bias, so that the sampling fraction was the same for all character states.

### BiSSE analyses

In a full BiSSE model, six parameters were estimated: the speciation rates for states 0 and 1 (*λ*_0_, *λ*_1_), the extinction rates for states 0 and 1 (*μ*_0_, *μ*_1_), and the transition rates between states 0 and 1 (*q*_01_, *q*_10_). For the self-compatibility character, we followed previous analyses (Goldberg et al., 2010; Freyman & Höhna, 2019) and constrained the transition rate from state 0 (SI) to state 1 (SC) to be 0 (i.e., *q*_10_ = 0). This constraint reflects the theoretical prediction and observation that the breakdown of self-incompatibility is rarely reversible (Igic et al., 2008; Barrett, 2013). When the character is pollination system or species-level outcrossing rate, transitions between state 0 and state 1 may be reversible. Therefore, we implemented both the full BiSSE model and a BiSSE model that assumes no transition from selfing (state 1) to outcrossing (state 0). If the two models produced qualitatively different results for a genus (see Table S3), the genus was considered sensitive to model assumptions and was dropped from the BiSSE MCMC analysis.

For each tree, we implemented BiSSE models using Markov chain Monte Carlo (MCMC) sampling (FitzJohn et al., 2009) to obtain the posterior probability distributions of each parameter. MCMC chains were started at the maximum likelihood estimates of the model parameters and were run for 2500 steps with the initial 20% steps discarded as the burn-in stage. For each genus, we merged the post-burn-in MCMC steps for all 100 bootstrap trees, resulting in a total of 200000 MCMC steps to account for phylogenetic uncertainty. Following Sabath et al. (2015), We calculated the proportion of MCMC steps in which the posterior diversification rates were higher for outcrossing lineages than for selfing lineages, denoted by *P*(*r*_0_>*r*_1_). Similar quantities were calculated for speciation rates and extinction rates, denoted by *P*(*λ*_0_>*λ*_1_) and *P*(*μ*_0_>*μ*_1_), respectively. These values should not significantly differ from 0.5 if the rate is state-independent.

For each rate parameter, the prior distribution used in the MCMC sampling was assumed to be exponential, with the mean being *fp*_0_. Here *p*_0_ is the initial rate estimate from the function “starting.point.bisse” in the *diversitree* package, and *f* = 1, 2, 4 is a parameter that we varied to examine whether the MCMC results were sensitive to the prior distributions (Sabath et al., 2015). Genera were considered as unrobust and were excluded from the BiSSE analyses if the *P*(*r*_0_ > *r*_1_) values obtained under the three values of *f* differed by > 30% (see Table S3). In the main text, we report the results obtained using *f* = 2.

To assess the significance of mating system on diversification across genera, we combined *P*(*r*_0_>*r*_1_) values of robust genera. Following Sabath et al (2015), we used a one-sample Wilcoxon rank-sum test. Under the null hypothesis of sate-independent diversification, the mean value of *P*(*r*_0_>*r*_1_) is 0.5, as diversification rates in state 0 should be higher than in state 1 for half of the MCMC steps. A similar test was conducted to evaluate the effects of mating systems on speciation rates, using *P*(*λ*_0_>*λ*_1_) values.

### MuSSE analyses

Our MuSSE analyses on the combination of self-compatibility and life form states were similar to the BiSSE analyses described above. However, the MuSSE model included four possible states: SI annual/short-lived (state 1), SI perennial (state 2), SC short-lived (state 3), and SC perennial (state 4). As in the BiSSE analyses, we assumed no transitions from SC to SI states. Transitions between life form states were reversible, as suggested from previous studies (Soltis et al., 2013). We also assumed that simultaneous transitions between self-compatibility and life form (e.g., from SI annual/short-lived to SC perennial) did not happen. Under these constraints, the MuSSE model estimated 14 parameters: the speciation rates for each state (*λ*_1_, *λ*_2_, *λ*_3_, *λ*_4_), the extinction rates for each state (*μ*_1_, *μ*_2_, *μ*_3_, *μ*_4_), transition rates in self-compatibility (*q*_13_, *q*_24_), and transition rates in life form (*q*_12_, *q*_21_, *q*_34_, *q*_43_). MCMC sampling was run for 10000 steps, with the initial 25% steps discarded as burn-in.

### HiSSE analyses

For each genus, we fitted six SSE models from the *hisse* package (Beaulieu & O’Meara, 2016) to the ML tree and 100 bootstrap trees. Three models included only two mating system states without hidden states, denoted as BiSSE. The other three models, labeled HiSSE, incorporated two hidden states, A and B, resulting in four possible states: 0A, 1A, 0B, and 1B. In all HiSSE models, we assumed no simultaneous transitions at both the mating system states and hidden states (e.g., no transition from 0A to 1B). For both model categories, we considered three types of model setups: a state-independent diversification model (BiSSE null, HiSSE CID2), a state-dependent diversification model with no backward transitions from state 1 (selfing) to state 0 (outcrossing) (BiSSE irr, HiSSE irr), and a state-dependent diversification model with reversible transitions between states 0 and 1 (BiSSE rev, HiSSE rev). It should be emphasized that the BiSSE models from the *hisse* package here differ from those in the *diversitree* package used in our MCMC analyses, as the model setups are not the same.

For each tree, we selected the best-fitting model from the six models based on the lowest AICc values (rather than AIC values) to account for the small sample size. For each genus, we summarized the best-fitting models across 100 bootstrap trees, and the best-fitted model for the genus was determined based on the models with highest proportion. For all genera, we found this model was the same as the best-fitting model identified from the ML tree.

When the model incorporating hidden states and state-dependency (HiSSE rev or HiSSE irr) provided the best fit for a genus, we assessed whether outcrossing consistently gave higher or lower diversification rates than selfing at both hidden states or not. To do this, we classified the rate estimates from each bootstrap tree into two scenarios: 1) diversification rates for state 0 (outcrossing) are either consistently higher or lower than for state 1 (selfing) in both hidden states (*r*_0*A*_>*r*_1*A*_ and *r*_0*B*_>*r*_0*B*_, or *r*_0*A*_ < *r*_1*A*_ and *r*_0*B*_ < *r*_0*B*_); 2) diversification rates for state 0 (outcrossing) are higher than for state 1 (selfing) in one hidden state but lower in the other hidden state (*r*_0*A*_>*r*_1*A*_ and *r*_0*B*_ < *r*_1*B*_, or *r*_0*A*_ < *r*_1*A*_ and *r*_0*B*_>*r*_1*B*_). We recorded the occurrence of the two scenarios in 100 bootstrap trees.

## Results

### Data description

After filtering for suitable genera, our diversification analyses include 27 genera, summarized in Table S2. Among these, 21 herbaceous genera are from the self-incompatibility dataset, which includes 9 genera in Asteraceae, 4 in Brassicaceae, 4 in Solanaceae, 2 in Bromeliaceae, 1 in Onagraceae, and 1 in Fabaceae. Four genera in Polemoniaceae are used based on the pollination system dataset. The outcrossing rate dataset provides 2 gymnosperm genera (*Picea, Pinus*) suitable for analyses. In *Oenothera*, some SC species are functionally asexual due to a unique genetic system known as permanent translocation heterozygosity (PTH), which nearly suppresses meiotic recombination and segregation (Holsinger & Ellstrand, 1984). These PTH species are thus dropped from the phylogeny used for diversification analyses.

In total, the diversification analysis included 360 species in state 0 and 453 species in state 1, with 19 genera having sampling fractions greater than 10% (Fig. S1). Two genera, *Solanum* and *Oenothera*, are suitable for performing diversification analyses on the combination of self-compatibility and life form states. For *Solanum*, the MuSSE analysis includes no SI annual/short-lived species, 73 SI perennial species, 25 SC annual/short-lived species, and 105 SC perennial species. After excluding PTH species, the analysis in *Oenothera* includes 9 SI annual/short-lived species, 31 SI perennial species, 21 SC annual/short-lived species, and 11 SC perennial species.

### BiSSE analysis on mating system states

Out of the 27 genera, BiSSE MCMC samplings in four genera (*Gilia, Leptosiphon, Nicotiana, Picea*) show high sensitivity to the choice of prior distributions or transition reversibility in mating systems (Table S3). Dropping these genera, our BiSSE analyses include 23 genera, covering 316 species in state 0 and 331 species in state 1. Fig. 1 presents the distribution *P*(*r*_0_>*r*_1_) and *P*(*λ*_0_>*λ*_1_) values obtained from the 100 bootstrap trees for each genus (detailed values are summarized in Table S4).

**Figure 1.**
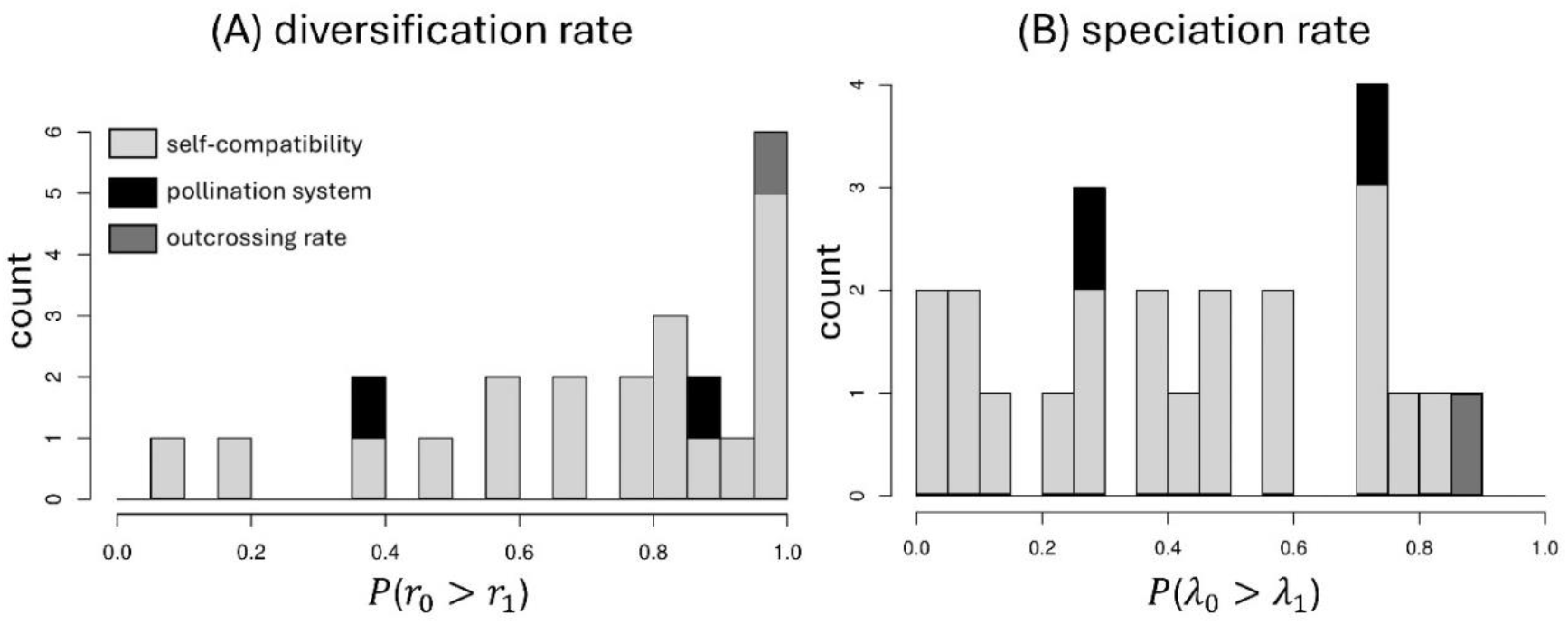
The proportion of the posterior probabilities from BiSSE MCMC sampling that supports higher diversification rates and speciation rates for state 0 (outcrossing) than state 1 (selfing), assuming no transition from state 1 to state 0. Colors represent different characters related to mating systems. The p-value from one-sample Wilcoxon rank-sum test is 0.002 for panel (A), and 0.111 for panel (B).

The results in Fig. 1(A) suggest that across genera, outcrossing (state 0) lineages have higher diversification rates than selfing (state 1) lineages, as *P*(*r*_0_>*r*_1_) values significantly deviate above 0.5 (*P*=0.002; one-sample Wilcoxon rank-sum test). In contrast, outcrossing lineages do not consistently speciate at higher or lower rates compared to selfing lineages, as *P*(*λ*_0_>*λ*_1_) values do not significantly deviate from 0.5 (*P*=0.286; one-sample Wilcoxon rank-sum test). The above results are robust to different values of *f* used in the prior distributions during MCMC sampling (Fig. S2).

Overall, the proportion of MCMC steps supporting higher diversification rates for outcrossing lineages, *P*(*r*_0_>*r*_1_), is positively correlated with that for speciation rates, *P*(*λ*_0_>*λ*_1_) (*r* = 0.45; *P*=0.032), while negatively correlated with the proportion supporting higher extinction rates, *P*(*μ*_0_>*μ*_1_) (*r* = -0.46; *P*=0.026). These results suggest that the higher diversification rates inferred for outcrossing lineages reflect both higher rates of speciation and lower rates of extinction. However, it should be noted that the inferred extinction rates may be unreliable (Rabosky, 2010).

Even if mating systems have no impact on diversification rates, genera with more outcrossing species tend to show higher *P*(*r*_0_>*r*_1_) values. However, we find no significant correlation between *P*(*r*_0_>*r*_1_) values and the fraction of outcrossing species included in the diversification analysis (*r*=0.15; *P*=0.500). This result suggests that the finding of higher *P*(*r*_0_>*r*_1_) values for outcrossing lineages is unlikely to be because outcrossing species happen to be overrepresented in our focal genera.

### Diversification analyses on the combination of self-compatibility and life form

We performed MuSSE MCMC analysis on the combination of self-compatibility and life form states in two genera, *Solanum* and *Oenothera*. For both genera, *P*(*r*_0_>*r*_1_) values from the BiSSE MCMC analysis are close to 1 (see Table S4), suggesting that SI lineages diversify at higher rates than SC lineages. Additional MuSSE analysis shows that within SC lineages, SC annual/short-lived lineages tend to diversify at much lower rates than SC perennial lineages.

In *Solanum*, SC annual/short-lived lineages exhibit much lower diversification rates than the other two states (Fig. 2A). This pattern appears to be driven by a combination of lower speciation rates (Fig. 2B) and higher extinction rates (Fig. S3). In contrast, SC perennial lineages display only slightly lower diversification rates than SI perennial lineages (Fig. 2A), which is likely driven by slightly higher extinction rates (Fig. S3) instead of difference in speciation rates (Fig. 2B). Additionally, transition rates from SI to SC are quite frequent among perennials (*q*_24_ in Fig. 2C). For life form evolution, the results show that the transition rates from SC perennials to SC annuals/short-lived species are rarely reversed (*q*_34_, *q*_43_ in Fig. 2C).

**Figure 2.**
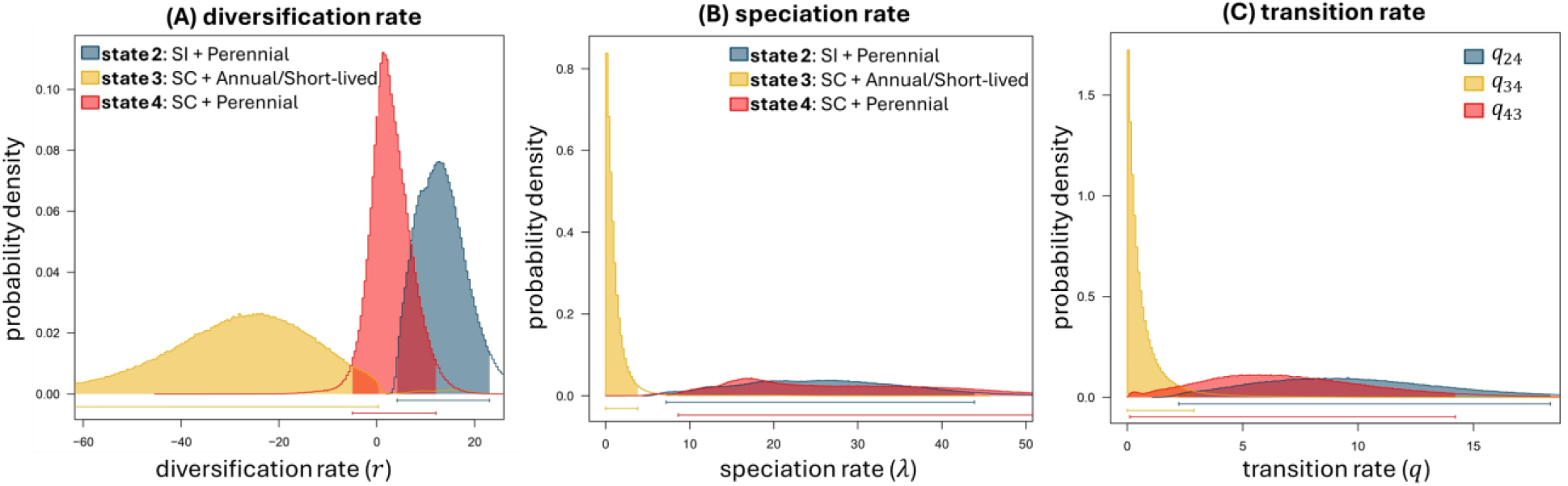
Posterior distribution of the diversification rates, speciation rates and transition rates for the combination of self-compatibility and life form states in *Solanum*. Results were obtained from MuSSE MCMC simulations on 100 bootstrap trees. Results are robust to different values of *f* used in the prior distribution (Fig. S3).

In *Oenothera*, SC lineages show lower diversification and speciation rates than SI lineages in both annual/short-lived and perennial states (Figs. 3A, 3B). However, the differences in diversification rates between SI and SC lineages are much more pronounced in the annual/short-lived state than in the perennial state (Fig. 3A). The low diversification rates in SC annual/short-lived lineages are suggested to be because extinction rates are greatly elevated (Fig. S4). However, among SI species, SI annual/short-lived lineages have higher diversification rates than SI perennials (Fig. 3A), mainly driven by higher speciation rates (Figs. 3B, S4). Transition rates from SI to SC are higher in annual/short-lived lineages than in perennial lineages (Fig. 3C). For transitions between life form states, the rate of transition from annuals/short-lived to perennials is much higher than in the reverse direction in SI lineages (compared *q*_12_, *q*_21_ in Fig. 3D). However, among SC lineages, the rate of transition toward perennials is greatly reduced, making the transition rates between life form states in both directions comparable (compared *q*_34_, *q*_43_ in Fig 3D).

**Figure 3.**
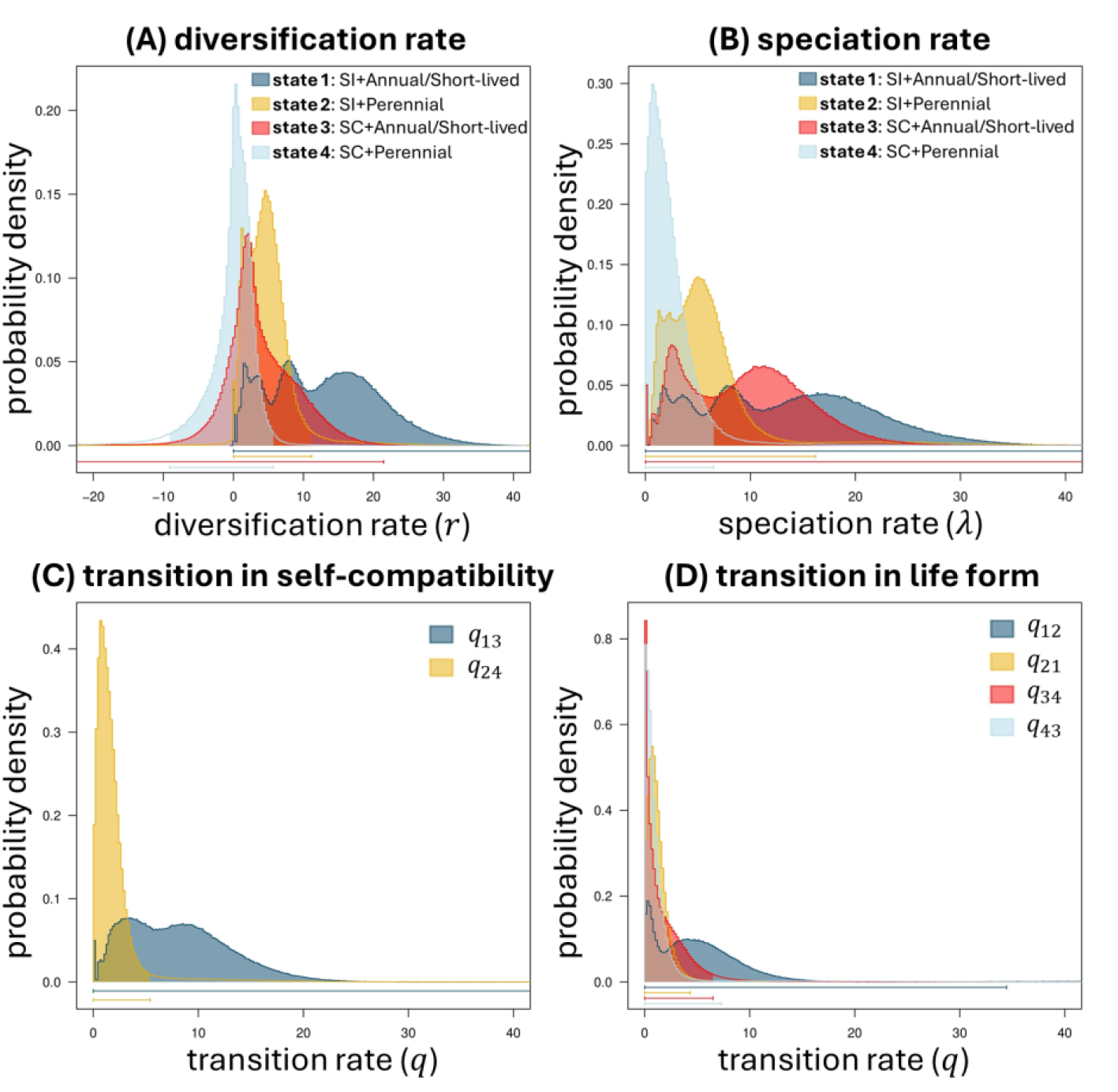
Posterior distribution of the diversification rates, speciation rates and transition rates for the combination of self-compatibility and life form states in *Oenothera*. Results were obtained from MuSSE MCMC simulations on 100 bootstrap trees. Results are robust to different values of *f* used in the prior distribution (Fig. S4).

### Diversification analysis with hidden states

Results from the MuSSE analysis indicate that the effects of selfing on lineage diversification may depend on the states of other characters. This conjecture is also supported by the HiSSE analysis. Out of the 27 focal genera, models incorporating hidden states are better fitted than models assuming no hidden states in 19 genera, as shown in Fig. 4A (detailed results are summarized in Table S5). Models assuming no backward transition from selfing to outcrossing (BiSSE irr, HiSSE irr) are best fitted in 12 genera (Fig. 4A). Models that allow reversible transitions between mating system states (BiSSE rev, HiSSE rev) are best fitting for the two genera from the outcrossing rate dataset (dark grey in Fig. 4A), suggesting that changes in the rate of selfing tends to be reversible. However, models allowing reversible transitions were also best fitted in 7 genera from the self-compatibility dataset. This may be due to overfitting and may not accurately reflect reality, as most of these genera belong to Asteraceae and Brassicaceae, where self-incompatibility systems are maintained by self-recognition mechanisms that should rarely allow transitions from SC back to SI (Fujii et al., 2016). Models incorporating hidden states and state-dependent diversification (HiSSE rev, HiSSE irr) are the best-fitting for 17 genera. However, no significant pattern is found regarding whether outcrossing lineages consistently diversify at higher or lower rates compared to selfing lineages across both hidden states (Fig. 4B).

**Figure 4.**
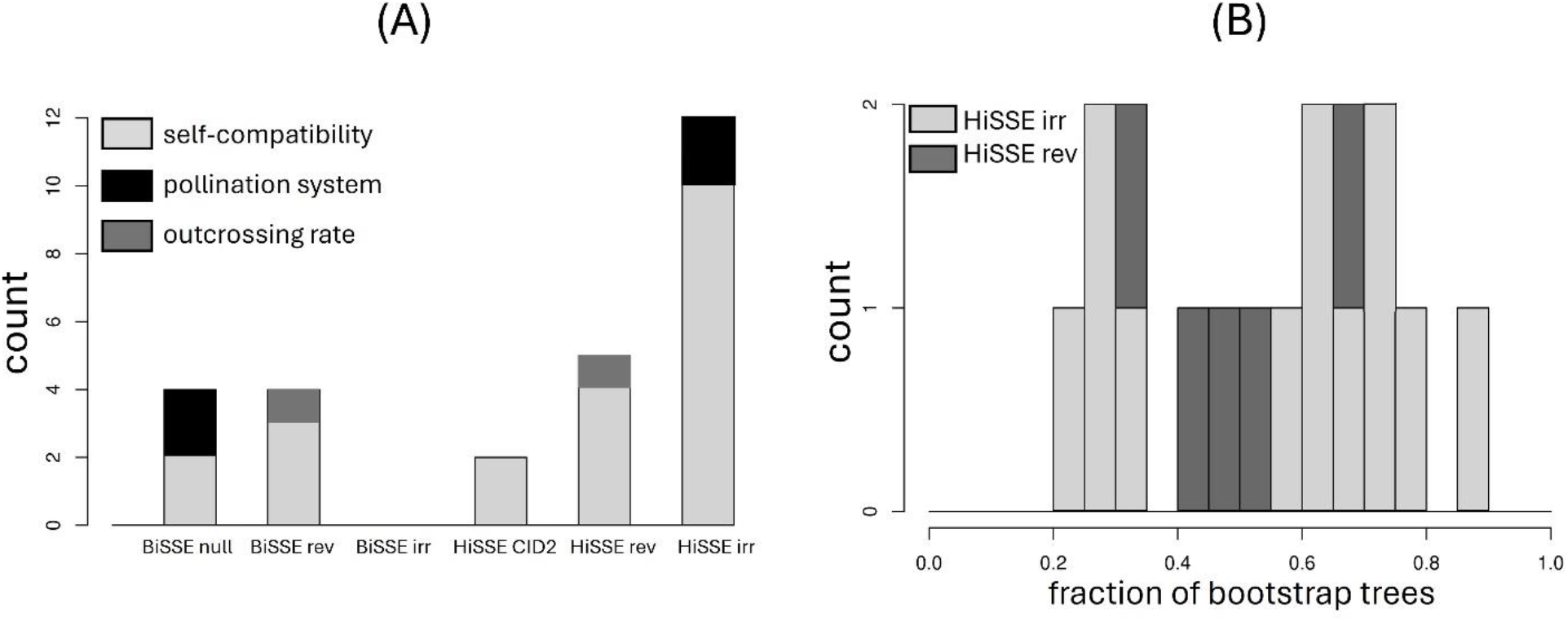
(A) Distribution of the best-fitting model across 27 genera. (B) The fraction of 100 bootstrap trees of each genus for which the estimated diversification rates from the *hisse* package for state 0 are consistently higher or lower than state 1 in both hidden states A and B. The distribution of the fractions does not significantly deviate from 0.5 in one-sample Wilcoxon rank-sum test, no matter whether the two HiSSE models were analyzed separately (*P* = 0.478 for HiSSE irr; *P* = 1.000 for HiSSE rev) or combined (*P* = 0.505). Detailed results for each genus are presented in Table S5.

## Discussion

The current study investigated the effects of mating system states on lineage diversification rates by performing phylogenetic-based diversification analyses to a dozen of genera. The results show that outcrossing lineages tend to have higher net diversification rates than selfing lineages, which is suggested to be attributed to both higher speciation rates and lower extinction rates. Nevertheless, mating systems do not exhibit consistent effects on speciation rates; selfing lineages display higher speciation rates in some genera while exhibiting lower rates in others. This result aligns with the mixed outcomes observed in previous analyses. This variability may stem from the fact that speciation events involve various mechanisms (Coyne & Orr, 2004), and selfing can have complex effects on these mechanisms, as discussed in the introduction, and reviewed by Marie-Orleach et al. (2024).

Most previous analyses were conducted at the family level, but the current genus-level analyses reveal that within the same family, the effects of mating systems on diversification rates can vary across genera. For instance, family-level analyses in Solanaceae reported lower diversification rates in self-compatible (SC) lineages compared to self-incompatible (SI) lineages (Goldberg et al., 2010). While similar results are observed in the genus *Solanum*, self-compatibility shows no significant effects on diversification in *Nicotiana*. Since one-third of the species included in Goldberg et al. (2010) are from *Solanum*, their results may be skewed by the patterns observed in this genus.

We propose that highly selfing lineages tend to have much higher extinction rates than mixed-mating lineages. Although we are not able to directly test this hypothesis due to insufficient data, we find SC annual/short-lived lineages exhibit much lower diversification rates than SC perennials in *Solanum*. Also, in *Oenothera*, diversification rate differences between SI and SC lineages are more prominent in annuals/short-lived than in perennials. These results indirectly support our hypothesis as annuals are more likely to be highly selfing than perennials (Aarssen, 2000; Ma et al., 2020). A direct test of this hypothesis would require a diversification analysis across a full range of selfing rates once outcrossing rates are measured in more species. Additionally, this hypothesis can be indirectly examined by conducting diversification analyses of quantitative traits related to selfing rates, such as anther-stigma distance and flower size (Thomann et al., 2013).

As explained in the introduction, the above results may help explain the enigma of the common occurrence of mixed-mating systems (Goodwillie et al., 2005). Additionally, our analyses indirectly support the model prediction that mixed-mating systems tend to be evolutionarily unstable (Lloyd, 1979; Lande & Schemske, 1985). Specifically, we find that transition rates from SC perennials, which tend to exhibit mixed mating, to SC annuals/short-lived plants, which are likely to be highly selfing, are higher than the reverse direction. However, we also find relatively high transition rates from SI perennials to SC perennials, which suggests that the prevalence of mixed mating systems may also be partly contributed by frequent transitions from outcrossing lineages.

The analyses incorporating life form states suggest that variation in diversification rates under different mating systems may often be partly driven by unmeasured factors associated with mating system transitions. Indeed, we find that SSE models incorporating hidden states are better fitted than those without hidden states for a majority of the studied genera. This may be partly because transitions from outcrossing to selfing are often triggered by environmental changes, such as colonization and pollen limitation (Busch & Schoen, 2008; Eckert et al., 2010; Pannell, 2016; Encinas-Viso et al., 2020). Nevertheless, how the impacts of selfing on diversification rates depend on hidden states is not consistent across genera. In some genera, selfing consistently promotes or inhibits diversification for both hidden states, but in other genera, selfing has opposite effects on diversification in different hidden states.

Our studies also support the model prediction that self-fertilization tends to promote evolution towards shorter lifespans (Lesaffre & Billiard, 2020). Among SC speciess in *Solanum*, transitions from perennials to annuals/short-lived lineages are found to be nearly unidirectional. Also, in *Oenothera*, although transition rates from perennials to annuals/short-lived are not elevated in SC lineages, the rate of transitions from annuals/short-lived to perennials is greatly reduced in SC lineages compared to SI lineages.

In short, the current analyses suggest that selfing lineages have lower diversification rates and higher extinction rates, which may be mainly contributed in highly selfing, instead of mixed-mating lineages. However, it should be emphasized that our analyses only reveal a correlation, instead of causation, between selfing and low diversification rates. In fact, the impacts of selfing on diversification are likely context-dependent, potentially interacting with other ecological or genetic factors.

## Acknowledgments

I would like to thank Stephen Wright and Mark Hibbins for comments and suggestions on the analyses and a previous version of the manuscript. K.X. was supported by the EEB Postdoctoral Fellowship offered by the Department of Ecology & Evolutionary Biology at the University of Toronto.

## Author contributions

K.X. conceived and designed the study, conducted the analysis and drafted the article.

## Data availability

Mating system character states, supermatrices and phylogenies of each genus are available at https://doi.org/10.5281/zenodo.13265323

