## Supplementary Figures and Tables for "Analysis of the effects of mating systems on lineage diversification across multiple genera"

**Supplementary Tables and Figures for "Analysis of the effects of mating systems on lineage diversification across multiple genera"**

**Table S1.** Outgroups used for phylogeny reconstruction of each genus and related references (see reference list at the end of this document). The outgroup of the four genera from the pollination system dataset is marked N/A since we directly used the family-level phylogenies from the reference.

| Genus | Outgroups | References |
|---|---|---|
| **Self-compatibility dataset** | | |
| *Aechmea* | *Bromelia balansae*, *Puya laxa*, *Puya raimondii* | Schulte et al., 2009 |
| *Bidens* | *Coreopsis grandiflora*, *Coreopsis lanceolata* | Kim et al., 1999 |
| *Brassica* | *Caulanthus amplexicaulis*, *Diplotaxis tenuifolia*, *Sisymbrium irio*, *Streptanthus drepanoides* | Warwick & Hall, 2009; Arias et al., 2014 |
| *Cardamine* | *Barbarea* vulgaris, *Camelina sativa*, *Lepidium sativum*, *Lepidium subulatum* | Huang et al., 2022 |
| *Carthamus* | *Centaurea cyanus*, *Centaurea maculosa* | Yang et al., 2023 |
| *Cirsium* | *Cynara cardunculus*, *Onopordum acanthium*, *Onopordum tauricum* | Slotta et al., 2012 |
| *Diplotaxis* | *Eruca vesicaria* | Franzke et al., 2017 |
| *Flaveria* | *Helenium autumnale*, *Haploesthes greggii*, *Sartwellia mexicana*, *Tanacetum parthenium* | McKown et al., 2005; Lyu et al., 2015 |
| *Layia* | *Holozonia filipes*, *Lagophylla ramosissima*, *Madia sativa* | Baldwin & Wessa, 2000; Baldwin, 2005 |
| *Lepidium* | *Cardamine hirsuta* | Cai & Ma, 2016; Ma et al., 2020 |
| *Microseris* | Agoseris glauca, *Nothocalais troximoides* | Lohwasser et al., 2004 |
| *Nicotiana* | *Petunia axillaris*, *Solanum agnewiorum*, *Symonanthus bancroftii* | Wang et al., 2022; Wang et al., 2023 |
| *Oenothera* | *Clarkia gracilis*, *Clarkia xantiana* | Overson et al., 2023 |
| *Petunia* | *Calibrachoa heterophylla*, *Calibrachoa pygmaea*, *Calibrachoa sendtneriana* | Reck-Kortmann et al., 2014 |
| *Physalis* | *Iochroma cyaneum*, *Vassobia breviflora*, *Witheringia solanacea* | del Pilar Zamora-Tavares et al., 2016 |
| *Senecio* | *Doronicum pardalianches*, *Jacobaea auricula*, *Jacobaea carniolica*, *Othonna capensis* | Pelser et al., 2007; Calvo et al., 2013; |
| *Simsia* | *Davilanthus davilae*, *Davilanthus hypargyreus*, *Tithonia diversifolia*, *Viguiera dentata* | Schilling & Panero, 2010 |
| *Solanum* | *Jaltomata bicolor*, *Jaltomata dentata*, *Jaltomata procumbens*, *Jaltomata sinuosa* | Goldberg et al., 2010; Gagnon et al., 2022 |
| *Symphyotrichum* | *Ampelaster carolinianus*, *Almutaster pauciflorus*, *Canadanthus modestus* | Vaezi & Brouillet, 2009 |
| *Tillandsia* | *Stigmatodon oliganthus*, *Vriesea carinata*, *Vriesea incurvata* | Gomes-da-Silva & Souza-Chies, 2018 |
| *Trifolium* | *Medicago sativa*, *Melilotus albus*, *Trigonella foenumgraecum* | Watson et al., 2000 |
| **Pollination system dataset** | | |
| *Cobaea* | N/A | Landis et al., 2018 |
| *Gilia* | N/A | Landis et al., 2018 |
| *Ipomopsis* | N/A | Landis et al., 2018 |
| *Leptosiphon* | N/A | Landis et al., 2018 |
| **Outcrossing rate dataset** | | |
| Picea | *Cathaya argyrophylla*, *Pinus cembra*, *Pinus sylvestris*, *Pseudotsuga menziesii* | Lockwood et al., 2013 |
| Pinus | *Cathaya argyrophylla*, *Cedrus deodara*, *Picea sitchensis*, *Pseudolarix amabilis*, *Pseudotsuga menziesii* | Gernandt et al., 2005; Palmé et al., 2009 |

**Table S2.** Genera included in the diversification analysis.

| Genus | Family | Species richness[a] | No. sp. tree[b] | State 0 | State 1 | No. clusters | Alignment Length | Missing site (%)[c] |
|---|---|---|---|---|---|---|---|---|
| **Self-compatibility dataset** | | | | | | | | |
| *Aechmea* | Bromeliaceae | 257 | 158 | 7 | 10 | 38 | 33323 | 78 |
| *Bidens* | Asteraceae | 246 | 88 | 5 | 30 | 25 | 20530 | 64 |
| *Brassica* | Brassicaceae | 46 | 27 | 13 | 3 | 75 | 54952 | 81 |
| *Cardamine* | Brassicaceae | 253 | 180 | 6 | 10 | 20 | 16044 | 82 |
| *Carthamus* | Asteraceae | 26 | 28 | 2 | 8 | 46 | 33372 | 67 |
| *Cirsium* | Asteraceae | 497 | 153 | 3 | 7 | 16 | 10869 | 55 |
| *Diplotaxis* | Brassicaceae | 40 | 29 | 13 | 3 | 26 | 17729 | 82 |
| *Flaveria* | Asteraceae | 21 | 20 | 16 | 3 | 43 | 40264 | 54 |
| *Layia* | Asteraceae | 16 | 10 | 7 | 3 | 18 | 14642 | 71 |
| *Lepidium* | Brassicaceae | 249 | 140 | 5 | 9 | 52 | 29997 | 86 |
| *Microseris* | Asteraceae | 44 | 15 | 4 | 5 | 12 | 8040 | 61 |
| *Nicotiana* | Solanaceae | 101 | 72 | 7 | 41 | 73 | 71676 | 81 |
| *Oenothera* | Onagraceae | 165 | 112 | 40 | 32 | 23 | 24604 | 76 |
| *Petunia* | Solanaceae | 16 | 17 | 8 | 3 | 38 | 30449 | 59 |
| *Physalis* | Solanaceae | 85 | 61 | 6 | 8 | 40 | 28310 | 72 |
| *Senecio* | Asteraceae | 1681 | 485 | 21 | 17 | 15 | 10944 | 70 |
| *Simsia* | Asteraceae | 30 | 22 | 14 | 3 | 7 | 3759 | 52 |
| *Solanum*[d] | Solanaceae | 1238 | 741 | 73 | 133 | 9 | 10908 | N/A |
| *Symphyotrichum* | Asteraceae | 104 | 64 | 5 | 4 | 39 | 22174 | 80 |
| *Tillandsia* | Bromeliaceae | 678 | 291 | 6 | 16 | 20 | 18552 | 71 |
| *Trifolium* | Fabaceae | 369 | 225 | 5 | 11 | 9 | 6382 | 53 |
| **Pollination system dataset** | | | | | | | | |
| *Cobaea*[e] | Polemoniaceae | 17 | N/A | 8 | 2 | N/A | N/A | N/A |
| *Gilia*[e] | Polemoniaceae | 39 | N/A | 10 | 21 | N/A | N/A | N/A |
| *Ipomopsis*[e] | Polemoniaceae | 27 | N/A | 15 | 4 | N/A | N/A | N/A |
| *Leptosiphon*[e] | Polemoniaceae | 33 | N/A | 7 | 4 | N/A | N/A | N/A |
| **Outcrossing rate dataset** | | | | | | | | |
| *Picea* | Pinaceae | 43 | 34 | 7 | 5 | 197 | 144673 | 75 |
| *Pinus* | Pinaceae | 134 | 116 | 34 | 7 | 362 | 174017 | 80 |

a. The number of accepted species per genus in WFO Plant List.
b. The number of species on the reconstructed phylogenetic tree.
c. Percentage of sites with missing data in the concatenated supermatrix.
d. Supermatrix used for phylogeny reconstruction of *Solanum* is directly taken from Gagnon et al. (2022).
e. ML trees and bootstrap trees of the genus are directly taken from the family-level phylogenies in Landis et al. (2018).

**Table S3.** Comparison of BiSSE models with and without transition from state 1 (selfing) to state 0 (outcrossing) when the mating-system-related character is pollination system or mean outcrossing rates. In the table, "BiSSE model with $q_{10} = 0$" stands for the BiSSE model from the *diversitree* package with no transition from state 1 to state 0. "full BiSSE model" represents the full BiSSE model without this constraint. The values reported in the table are the proportion of posterior distribution supporting higher diversification rates for state 0 than state 1, $P(r_0 > r_1)$, under different values of parameter $f$ used in the prior distribution during MCMC sampling.

| **Genus** | **Family** | **BiSSE model with $q_{10} = 0$** | | | **full BiSSE model** | | |
|---|---|---|---|---|---|---|---|
| | | $f = 1$ | $f = 2$ | $f = 4$ | $f = 1$ | $f = 2$ | $f = 4$ |
| **Pollination system dataset** | | | | | | | |
| *Cobaea* | Polemoniaceae | 0.40 | 0.38 | 0.36 | 0.51 | 0.38 | 0.27 |
| *Gilia* | Polemoniaceae | 1.00 | 1.00 | 1.00 | 0.06 | 0.06 | 0.06 |
| *Ipomopsis* | Polemoniaceae | 0.91 | 0.93 | 0.95 | 0.88 | 0.84 | 0.82 |
| *Leptosiphon* | Polemoniaceae | 0.51 | 0.54 | 0.58 | 0.54 | 0.36 | 0.27 |
| **Outcrossing rate dataset** | | | | | | | |
| *Picea* | Pinaceae | 0.96 | 0.97 | 0.98 | 0.45 | 0.39 | 0.38 |
| *Pinus* | Pinaceae | 0.71 | 0.78 | 0.86 | 0.97 | 0.96 | 0.90 |

**Table S4.** Proportion of posterior distributions from the BiSSE MCMC analysis supporting higher diversification rates, speciation rates, and extinction rates for state 0 than state 1. $f$ is the parameter used in the prior distributions during MCMC sampling, described in the main text. Results shown are from the BiSSE model that assumes no transitions from state 1 (selfing) to state 0 (outcrossing). Genera colored in red exhibit high sensitivity to the choice of prior distributions or reversibility in mating system states.

| | $P(r_0 > r_1)$ | | | $P(\lambda_0 > \lambda_1)$ | | | $P(\mu_0 > \mu_1)$ | | |
|---|---|---|---|---|---|---|---|---|---|
| | $f = 1$ | $f = 2$ | $f = 4$ | $f = 1$ | $f = 2$ | $f = 4$ | $f = 1$ | $f = 2$ | $f = 4$ |
| **Aechmea** | 0.66 | 0.77 | 0.87 | 0.49 | 0.58 | 0.63 | 0.13 | 0.12 | 0.11 |
| **Bidens** | 0.16 | 0.20 | 0.42 | 0.01 | 0.01 | 0.01 | 0.10 | 0.04 | 0.01 |
| **Brassica** | 0.80 | 0.84 | 0.87 | 0.72 | 0.72 | 0.71 | 0.45 | 0.43 | 0.40 |
| **Cardamine** | 0.66 | 0.76 | 0.88 | 0.46 | 0.47 | 0.44 | 0.14 | 0.11 | 0.08 |
| **Carthamus** | 0.74 | 0.87 | 0.95 | 0.57 | 0.71 | 0.80 | 0.04 | 0.04 | 0.03 |
| **Cirsium** | 0.49 | 0.57 | 0.66 | 0.29 | 0.36 | 0.40 | 0.15 | 0.11 | 0.09 |
| **Diplotaxis** | 0.85 | 0.86 | 0.89 | 0.40 | 0.26 | 0.15 | 0.02 | 0.02 | 0.02 |
| **Flaveria** | 0.95 | 0.96 | 0.96 | 0.68 | 0.57 | 0.45 | 0.00 | 0.00 | 0.00 |
| **Layia** | 0.78 | 0.81 | 0.84 | 0.81 | 0.85 | 0.88 | 0.71 | 0.71 | 0.71 |
| **Lepidium** | 0.55 | 0.67 | 0.78 | 0.31 | 0.41 | 0.49 | 0.10 | 0.08 | 0.06 |
| **Microseris** | 0.02 | 0.06 | 0.15 | 0.02 | 0.09 | 0.23 | 0.73 | 0.71 | 0.71 |
| **Nicotiana** | 0.23 | 0.52 | 0.72 | 0.13 | 0.26 | 0.35 | 0.28 | 0.25 | 0.22 |
| **Oenothera** | 0.95 | 0.99 | 1.00 | 0.72 | 0.76 | 0.80 | 0.17 | 0.15 | 0.15 |
| **Petunia** | 0.31 | 0.35 | 0.38 | 0.31 | 0.36 | 0.39 | 0.48 | 0.48 | 0.48 |
| **Physalis** | 0.72 | 0.84 | 0.91 | 0.64 | 0.72 | 0.75 | 0.33 | 0.32 | 0.31 |
| **Senecio** | 1.00 | 1.00 | 1.00 | 0.41 | 0.14 | 0.04 | 0.02 | 0.00 | 0.00 |
| **Simsia** | 0.57 | 0.59 | 0.62 | 0.48 | 0.45 | 0.44 | 0.24 | 0.23 | 0.21 |
| **Solanum** | 0.98 | 1.00 | 1.00 | 0.00 | 0.00 | 0.00 | 0.00 | 0.00 | 0.00 |
| **Symphyotrichum** | 0.41 | 0.47 | 0.58 | 0.26 | 0.28 | 0.32 | 0.10 | 0.10 | 0.09 |
| **Tillandsia** | 0.97 | 1.00 | 0.98 | 0.46 | 0.21 | 0.52 | 0.06 | 0.00 | 0.08 |
| **Trifolium** | 0.60 | 0.70 | 0.75 | 0.10 | 0.09 | 0.09 | 0.04 | 0.02 | 0.01 |
| **Cobaea** | 0.40 | 0.38 | 0.36 | 0.33 | 0.30 | 0.27 | 0.30 | 0.30 | 0.30 |
| **Gilia** | 0.99 | 1.00 | 1.00 | 0.98 | 0.99 | 0.99 | 0.00 | 0.00 | 0.00 |
| **Ipomopsis** | 0.91 | 0.93 | 0.95 | 0.76 | 0.72 | 0.65 | 0.00 | 0.00 | 0.00 |
| **Leptosiphon** | 0.51 | 0.54 | 0.58 | 0.44 | 0.45 | 0.45 | 0.18 | 0.18 | 0.18 |
| **Eucalyptus** | 0.97 | 0.97 | 0.97 | 0.51 | 0.56 | 0.60 | 0.02 | 0.02 | 0.02 |
| **Picea** | 0.71 | 0.78 | 0.86 | 0.41 | 0.40 | 0.38 | 0.07 | 0.05 | 0.03 |
| **Pinus** | 0.96 | 0.97 | 0.98 | 0.88 | 0.85 | 0.80 | 0.16 | 0.13 | 0.11 |

**Table S5**. Best-fitting model for each genus inferred from *hisse* package based on the lowest AIC values or AICc values. For genera with "HiSSE irr" or "HiSSE rev" being the best-fitting model, the last three columns give the fraction of 100 bootstrap trees with rate estimations falling in three categories: diversification rates for state 0 are higher than for state 1 at both hidden states ($r_{0A} > r_{1B}$, $r_{0B} > r_{1B}$), diversification rates for state 0 are lower than for state 1 at both hidden states ($r_{0A} < r_{1B}$, $r_{0B} < r_{1B}$), and diversification rates for state 0 are higher than state1 at one hidden state while lower at the other ($r_{0A} < r_{1B}$, $r_{0B} > r_{1B}$ or $r_{0A} > r_{1B}$, $r_{0B} < r_{1B}$).

| **Genus** | **Family** | **Best-fitting model based on AIC** | **Best-fitting model based on AICc** | $r_{0A} > r_{1B}$ $r_{0B} > r_{1B}$ | $r_{0A} < r_{1B}$ $r_{0B} < r_{1B}$ | $r_{0A} < r_{1B}$ $r_{0B} > r_{1B}$ **OR** $r_{0A} > r_{1B}$ $r_{0B} < r_{1B}$ |
|---|---|---|---|---|---|---|
| **Self-compatibility** | | | | | | |
| *Aechmea* | Bromeliaceae | BiSSE null | BiSSE null | | | |
| *Bidens* | Asteraceae | BiSSE rev | BiSSE rev | | | |
| *Brassica* | Brassicaceae | HiSSE irr | HiSSE rev | 25 | 17 | 58 |
| *Cardamine* | Brassicaceae | BiSSE null | HiSSE rev | 5 | 50 | 31 |
| *Carthamus* | Asteraceae | BiSSE null | HiSSE irr | 76 | 1 | 23 |
| *Cirsium* | Asteraceae | BiSSE null | HiSSE irr | 55 | 3 | 41 |
| *Diplotaxis* | Brassicaceae | BiSSE rev | HiSSE rev | 47 | 6 | 47 |
| *Flaveria* | Asteraceae | BiSSE rev | BiSSE rev | | | |
| *Layia* | Asteraceae | BiSSE null | HiSSE irr | 60 | 0 | 38 |
| *Lepidium* | Brassicaceae | HiSSE CID2 | HiSSE irr | 84 | 2 | 12 |
| *Microseris* | Asteraceae | BiSSE null | HiSSE irr | 5 | 20 | 75 |
| *Nicotiana* | Solanaceae | HiSSE CID2 | HiSSE CID2 | | | |
| *Oenothera* | Onagraceae | HiSSE irr | HiSSE irr | 71 | 1 | 28 |
| *Petunia* | Solanaceae | BiSSE null | HiSSE irr | 19 | 7 | 72 |
| *Physalis* | Solanaceae | BiSSE rev | HiSSE irr | 62 | 1 | 36 |
| *Senecio* | Asteraceae | HiSSE CID2 | HiSSE CID2 | | | |
| *Simsia* | Asteraceae | BiSSE rev | BiSSE rev | | | |
| *Solanum* | Solanaceae | HiSSE rev | HiSSE rev | 11 | 22 | 67 |
| *Symphyotrichum* | Asteraceae | BiSSE null | HiSSE irr | 25 | 4 | 71 |
| *Tillandsia* | Bromeliaceae | BiSSE null | BiSSE null | | | |
| *Trifolium* | Fabaceae | BiSSE irr | HiSSE rev | 9 | 39 | 52 |
| **Pollination system dataset** | | | | | | |
| *Cobaea* | Polemoniaceae | BiSSE null | HiSSE irr | 16 | 17 | 67 |
| *Gilia* | Polemoniaceae | BiSSE null | BiSSE null | | | |
| *Ipomopsis* | Polemoniaceae | BiSSE null | BiSSE null | | | |
| *Leptosiphon* | Polemoniaceae | BiSSE null | HiSSE irr | 66 | 0 | 34 |
| **Outcrossing rate dataset** | | | | | | |
| *Pinus* | Pinaceae | BiSSE null | BiSSE rev | | | |
| *Picea* | Pinaceae | BiSSE irr | HiSSE irr | 71 | 0 | 29 |

**Supplementary Figures**

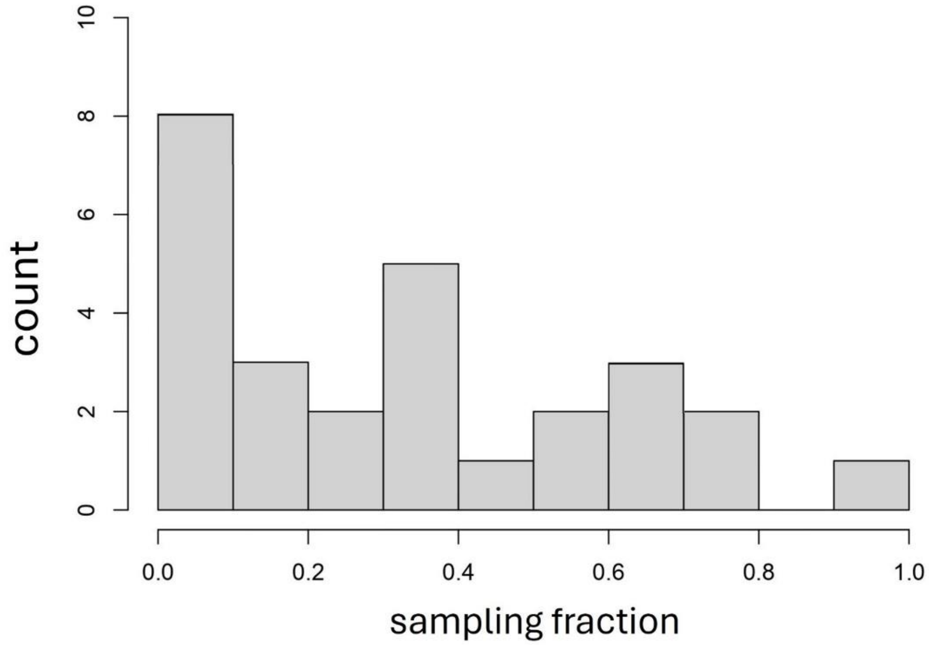


**Figure S1.** Distribution of the fraction of species included in the diversification analysis out of the number of accepted species in a genus, calculated from Table S1.

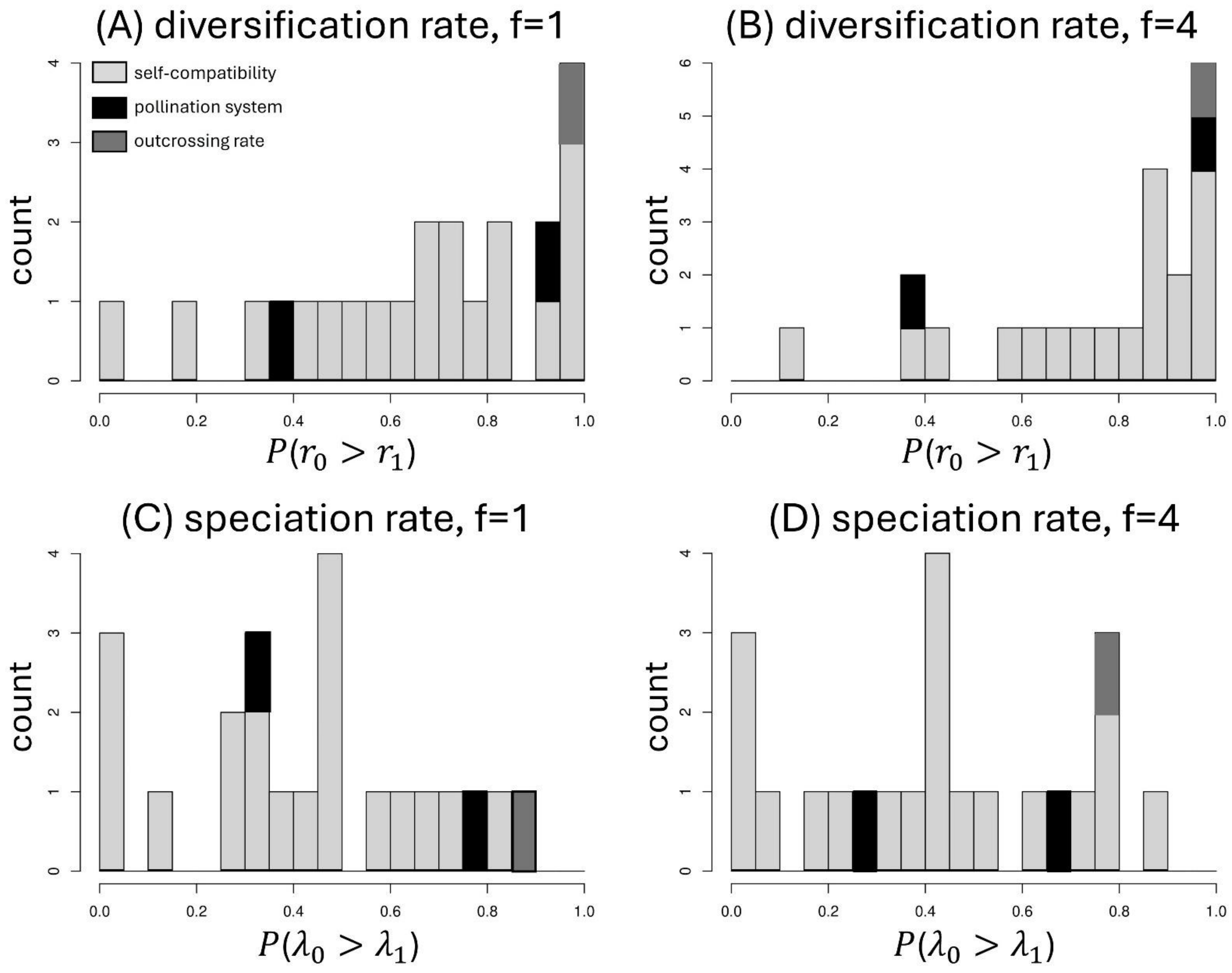


**Figure S2.** Proportion of the posterior probability distribution from BiSSE MCMC simulations that supports higher diversification rates and speciation rates for state 0 than state 1, assuming no transition from state 1 to state 0. Results are robust to different values of $f$ used in the prior distribution of the MCMC sampling. The p-value based on one-sample Wilcoxon rank-sum test is $P$ = 0.019 for panel (A), $P$ = 0.0001 for panel (B), $P$ = 0.222 for panel (C), and $P$ = 0.276 for panel (D).

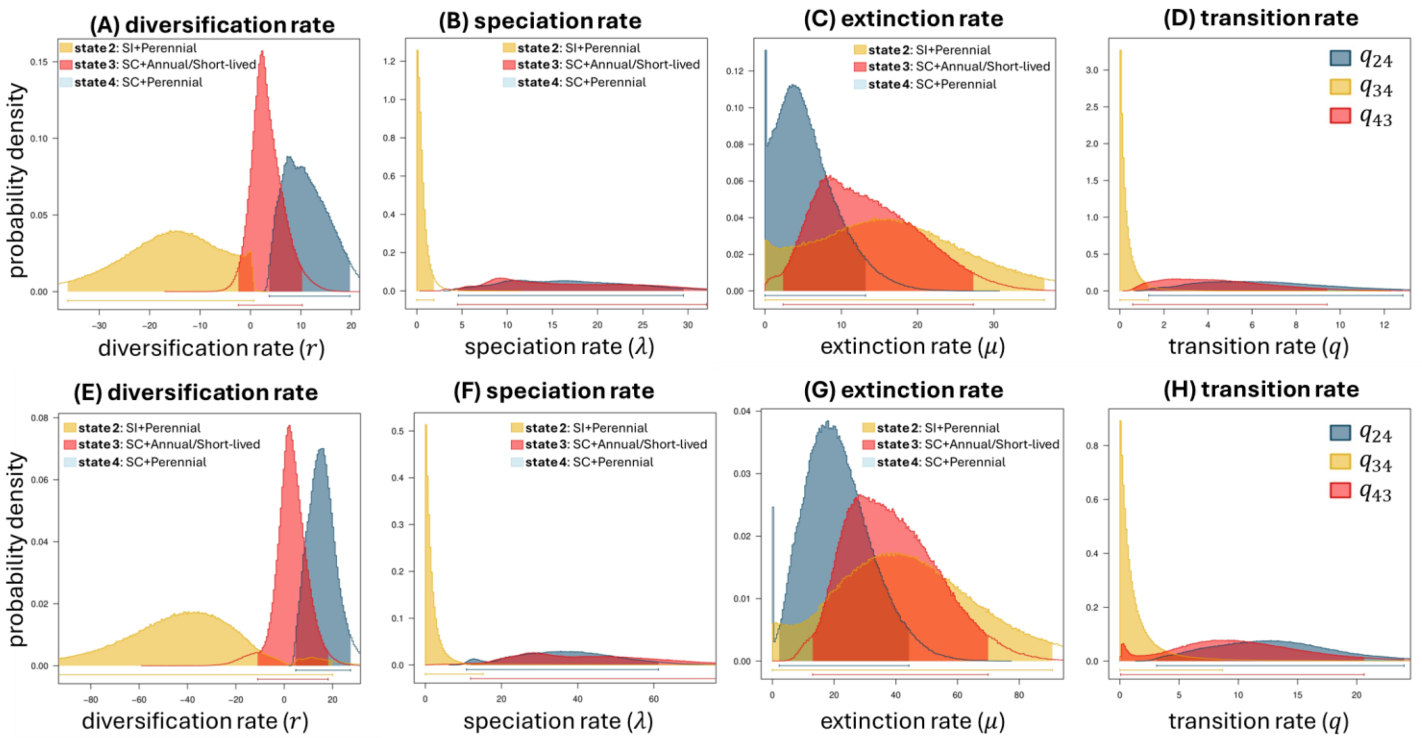


**Figure S3.** Posterior distribution of the net diversification rates, speciation rates, extinction rates, and transition rates obtained from MuSSE MCMC simulations on 100 bootstrap trees of *Solanum*. The parameter $f$ used in the prior distribution is $f = 1$ for upper panels and $f = 4$ for bottom panels.

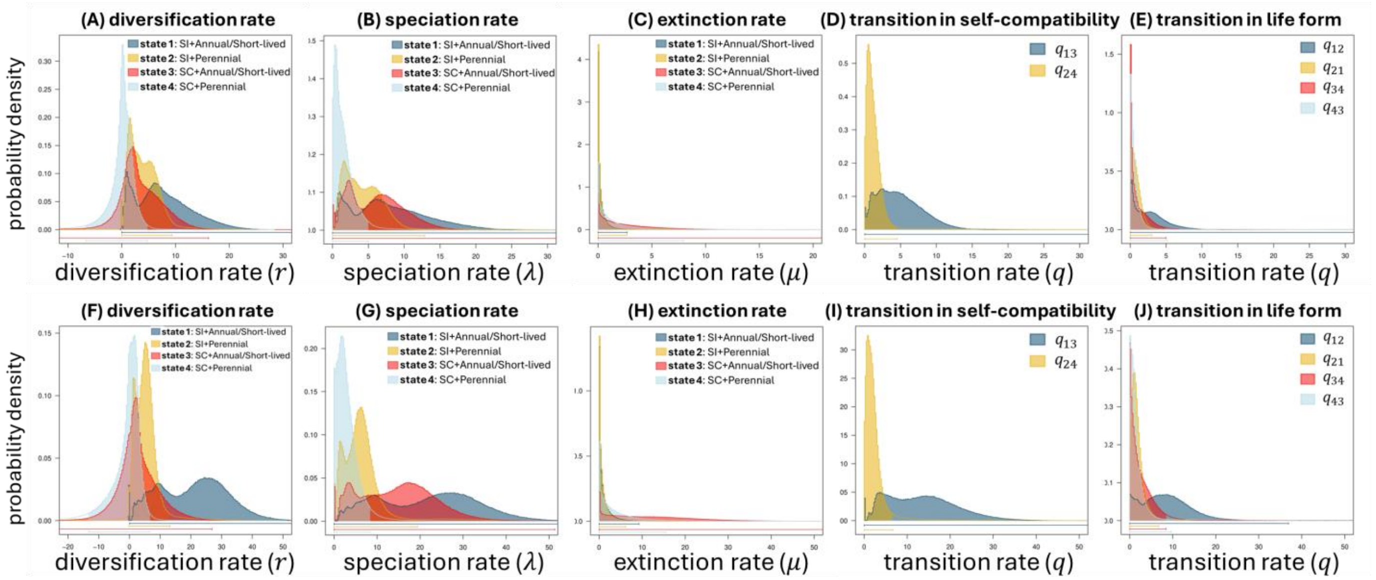


**Figure S4.** Posterior distribution of the net diversification rates, speciation rates, and transition rates obtained from MuSSE MCMC simulations on 100 bootstrap trees of *Oenothera*. The parameter $f$ used in the prior distribution is $f = 1$ for upper panels and $f = 4$ for bottom panels.
